# Variation in the Distribution of Large-scale Spatiotemporal Patterns of Activity Across Brain States

**DOI:** 10.1101/2024.04.26.591295

**Authors:** Lisa Meyer-Baese, Nmachi Anumba, T Bolt, L Daley, TJ LaGrow, Xiaodi Zhang, Nan Xu, Wen-Ju Pan, E Schumacher, Shella Keilholz

## Abstract

A few large-scale spatiotemporal patterns of brain activity (quasiperiodic patterns or QPPs) account for most of the spatial structure observed in resting state functional magnetic resonance imaging (rs-fMRI). The QPPs capture well-known features such as the evolution of the global signal and the alternating dominance of the default mode and task positive networks. These widespread patterns of activity have plausible ties to neuromodulatory input that mediates changes in nonlocalized processes, including arousal and attention. To determine whether QPPs exhibit variations across brain conditions, the relative magnitude and distribution of the three strongest QPPs were examined in two scenarios. First, in data from the Human Connectome Project, the relative incidence and magnitude of the QPPs was examined over the course of the scan, under the hypothesis that increasing drowsiness would shift the expression of the QPPs over time. Second, using rs-fMRI in rats obtained with a novel approach that minimizes noise, the relative incidence and magnitude of the QPPs was examined under three different anesthetic conditions expected to create distinct types of brain activity. The results indicate that both the distribution of QPPs and their magnitude changes with brain state, evidence of the sensitivity of these large-scale patterns to widespread changes linked to alterations in brain conditions.

## Introduction

Resting state functional magnetic resonance imaging (rs-fMRI) captures the spatiotemporal organization of the intrinsic activity of the brain and is a powerful translational tool for understanding normal and pathological brain function in humans and animals^1–7^. Recent work in healthy human subjects has shown that most of the features of rs-fMRI data, including functional connectivity, coactivation patterns, functional connectivity gradients, and modularity, can be explained by three repeated spatiotemporal patterns that cover the entire brain^8,9^. These patterns, sometimes called quasiperiodic patterns or QPPs^10–13^, capture cyclical activation and deactivation of brain areas and propagation of activity as areas transition between activated and deactivated phases. In humans, the first QPP describes the spatiotemporal evolution of the global signal, which contains contributions from both neural activity and widespread noise^14–18^. QPP2 captures the alternating activation and deactivation of the default mode network (DMN) and task positive network (TPN), which has been implicated in variability in task performance^19–21^. In the third QPP, somatomotor areas and visual areas activate and deactivate with opposite phases.

The spatial structure of the intrinsic activity captured by QPPs is remarkably similar across individuals and even across species^4,12^. While changes in functional connectivity, for example, are observed during deep sleep and anesthesia^22–24^, they are typically subtle. This stereotyped intrinsic activity interacts with tasks or stimuli and accounts for a portion of intraindividual variability^5,20,21,25,26^. Researchers increasingly use the term “brain state” to refer to commonly repeated configurations of brain activity. Because these states are dominated by intrinsic activity, and because intrinsic activity consists mostly of a few QPPs, we hypothesize that the relative expression of these spatiotemporal patterns defines brain states and reflects nonlocalized changes in brain activity related to phenomena such as arousal, focus, or emotion. For example, the global signal has previously been linked to arousal levels, with its amplitude increasing as arousal decreases^18,27,28^. Given the close links between QPP1 and the global signal, and between global signal and arousal, it seems likely that QPP1 might increase as arousal decreases. Preliminary data suggest that neuromodulatory nuclei play a role in driving these patterns of activity^29^, consistent with a potential link to changing conditions in the brain.

QPP2 also has potential links to arousal and/or attention. Prior studies of spatiotemporal patterns have typically utilized global signal regression, which suppresses QPP1 and makes QPP2 the primary pattern^5,13,25,30^. QPP2 is closely related to anticorrelated activity in the DMN and TPN, which has long been linked to task performance. In both humans and rats, QPP2 interacts with incoming stimuli and influences the resulting activation^5,26^. The spatial extent of the activity can be modulated by task performance in humans^13^, and the magnitude of the difference between activated and deactivated brain regions is greater when subjects are ‘in the zone’ on a finger tapping task^25^, but reduced in patients with ADHD compared to controls^30^. These converging findings suggest that QPP2 also has ties to attention and/or arousal, but because QPP1 was not examined, it is difficult to compare the relative sensitivity of the two patterns. Little work to date has examined QPP3.

In this study, we aim to understand how alterations in brain state affect the first three QPPs, taking a two-pronged approach. First, we perform further analysis of the human rs-fMRI data examined in ^8^, to determine whether the magnitude and/or relative incidence of different QPPs is constant or variable over the course of a scan where subjects may experience fluctuating arousal levels. We then examine rs-fMRI data acquired in rodents, where brain state is manipulated using three anesthetic conditions with different mechanisms of action and different effects on neural activity and vascular tone. Results from both species suggest that the strength and relative distribution of QPPs provide information about changes in brain state.

## Methods

### Overview

Resting state fMRI from human subjects was obtained from the Human Connectome Project^31^ and used to show that large-scale spatiotemporal patterns explain most of the spatial structure of functional connectivity in ^8^. We built on the original analysis of this data to examine the distribution of QPPs over the course of the scan. For rodents, all experimental protocols were approved by the Institutional Animal Care and Use Committee at Emory University. We first obtained rs-fMRI data from a small group of animals during free breathing or mechanical ventilation phase-locked to MRI acquisition to evaluate respiratory noise and motion, which can both contribute to large scale patterns of activity such as the global signal^16^. We then examined rs-fMRI data from mechanically-ventilated rats under three anesthetic conditions (reported previously in ^14^) to examine the spatiotemporal organization of intrinsic brain activity under conditions of minimal noise.

### Human Connectome Project data and analysis

We examined the magnitude of the time courses of the first three QPPs in the same 50 unrelated subjects (21 males) from the Human Connectome Project dataset used in the Bolt et al. paper (GE-EPI; TR 720 ms, 1200 volumes)^8^. To briefly summarize the prior analysis, the data was in surface based format (cifti) and had been denoised using HCP’s ICA-based algorithm. Spatial smoothing and bandpass filtering (0.01-0.1 Hz) were applied prior to further analysis. QPPs were extracted using the complex PCA approach described in ^8^ with code available at (https://github.com/tsb46/complex_pca). Ten components were obtained and varimax rotation was performed. The explained variance for each component and the magnitude and phase of each component at each time point were obtained. Based on the explained variance, three QPPs were retained for the remaining analysis.

To build upon the prior analysis and investigate the occurrence of different QPPs over the course of the scan, each time point was assigned a value (e.g., 1, 2, 3) that corresponded to the magnitude of the strongest QPP at that time point. This is a simplistic definition of brain state, but provides clear visualization of changes in the dominant QPP over time. Carpet plots of the state occurrence for each time point from each participant were created and sorted across time and across subject. The relative incidence of each state was calculated for each subject, and substantial intersubject variability was observed. To further explore this variability, subjects were clustered into two groups based on the mean and standard deviation of each component over the course of the scan using the k-means algorithm (Euclidean distance, 50 repetitions; k=2 chosen based on Silhouette criteria). The resulting groups were sorted across time. It was observed that the distribution of QPPs changed over the course of the scan, so we calculated the mean and standard deviation of the magnitude of each QPP for the first 200 and last 200 time points separately to detect any changes in QPP strength over the course of the scan. A paired t-test was applied for each QPP to identify significant changes.

Note that Bolt et al. used complex principal component analysis (cPCA) to extract the spatiotemporal patterns and that they are referred to as cPCs in ^8^. For consistency with prior work, we use the term QPP in this manuscript. The patterns extracted by the original QPP pattern-finding algorithm^9,11,12^ and the cPCA approach are highly similar^8^.

### Rodent data acquisition

One of the advantages of the rodent model is that motion-related noise is minimal relative to human studies. The rats are anesthetized and head-fixed, nearly eliminating motion of the head. However, the motion of the chest during respiration can affect the magnetic field at the brain, adding respiratory noise, and this effect is particularly strong in rodents, where the field strength is high and the chest is in close proximity to the head^32^. To minimize effects of differences in non-neural processes such as respiration across anesthetic conditions, we implemented a protocol where the rats are paralyzed, intubated and mechanically ventilated at frequency locked to image acquisition, ensuring that each slice is acquired at the same respiratory phase for each imaging volume. As shown in the Supplemental Material, this method reduces unwanted variability in rs-fMRI compared to the same rats breathing freely. The protocol was then applied in rats under three different anesthetic conditions to characterize the contributions of QPPs under each condition.

Acquisition of the data used for this analysis was reported previously^14^. Briefly, eight male Sprague Dawley rats (299g – 339g, Charles River) were intubated and ventilated at approximately 1 Hz. An infusion line was inserted subcutaneously to administer the paralytic pancuronium at a rate of 1.5 mg/kg/hr for the duration of the scan. For the comparison of different anesthetic conditions, rats were scanned consecutively under three protocols: 1.5% isoflurane (ISO), dexmedetomidine (DMED) only, and a combination of dexmedetomidine with a low dose isoflurane (ISODMED). These anesthetics have different effects on brain activity, and so would be expected to produce different brain states. Animals were first scanned for an average of 35 minutes under ISO. Rats were then subcutaneously injected with a 0.025 mg/kg bolus of DMED, taken off ISO five minutes later, and then switched to a 0.05 mg/kg/hr subcutaneous infusion of DMED at 10 minutes post-bolus. The animals were then scanned again for an average of 61 minutes. Afterwards, the animals were introduced to a low dose of ISO at 0.5% in combination with the 0.05 mg/kg/hr subcutaneous infusion of DMED. The animals were then scanned for a final time for 49 minutes on average. A table with the exact timing of anesthetic introduction and image acquisition for each rat is included in the supporting information of ^14^.

All rodent imaging was performed on a 9.4T/20 cm horizontal bore small animal MRI system with a homemade transmit/receive surface coil ∼2 cm in diameter. Anatomical scans were obtained using a T2-weighted RARE sequence (TR = 3500 ms, TE = 11 ms, 24 axial slices, 0.5 mm^3^ isotropic voxels). Resting state fMRI scans were acquired using a gradient-echo echo-planar imaging (EPI) sequence with field of view (FOV) 35mm x 35 mm, matrix size 70x70, and 24 axial slices for whole-brain coverage, resulting in isotropic voxels of 500 microns (TE = 15 ms, TR = 2000 ms) with partial Fourier encoding (encoding factor 1.4) to reduce the length of the echo train. A 3-volume reversed blip EPI image with the same parameters was acquired before each longer functional scan for TOPUP correction^33,34^. EPI scans included saturation bands to minimize signal from frontal and ventral regions outside the brain and were preceded by 10 dummy scans to ensure the signal reached steady state.

All rs-fMRI scans for the mechanically-ventilated rats were phase-locked, meaning that the time between subsequent images was set to a multiple of the animal’s respiratory rate so that each image was acquired during the same phase of the respiratory cycle^14^ (**Fig. 1**). This practice was adopted here to limit any effects of motion that could arise due to movement of the chest cavity and volumes being imaged at different points in the respiratory cycle. Images were acquired at a frequency of every other breath, or 2 Hz (TR = 2000 ms).

**Fig. 1.**
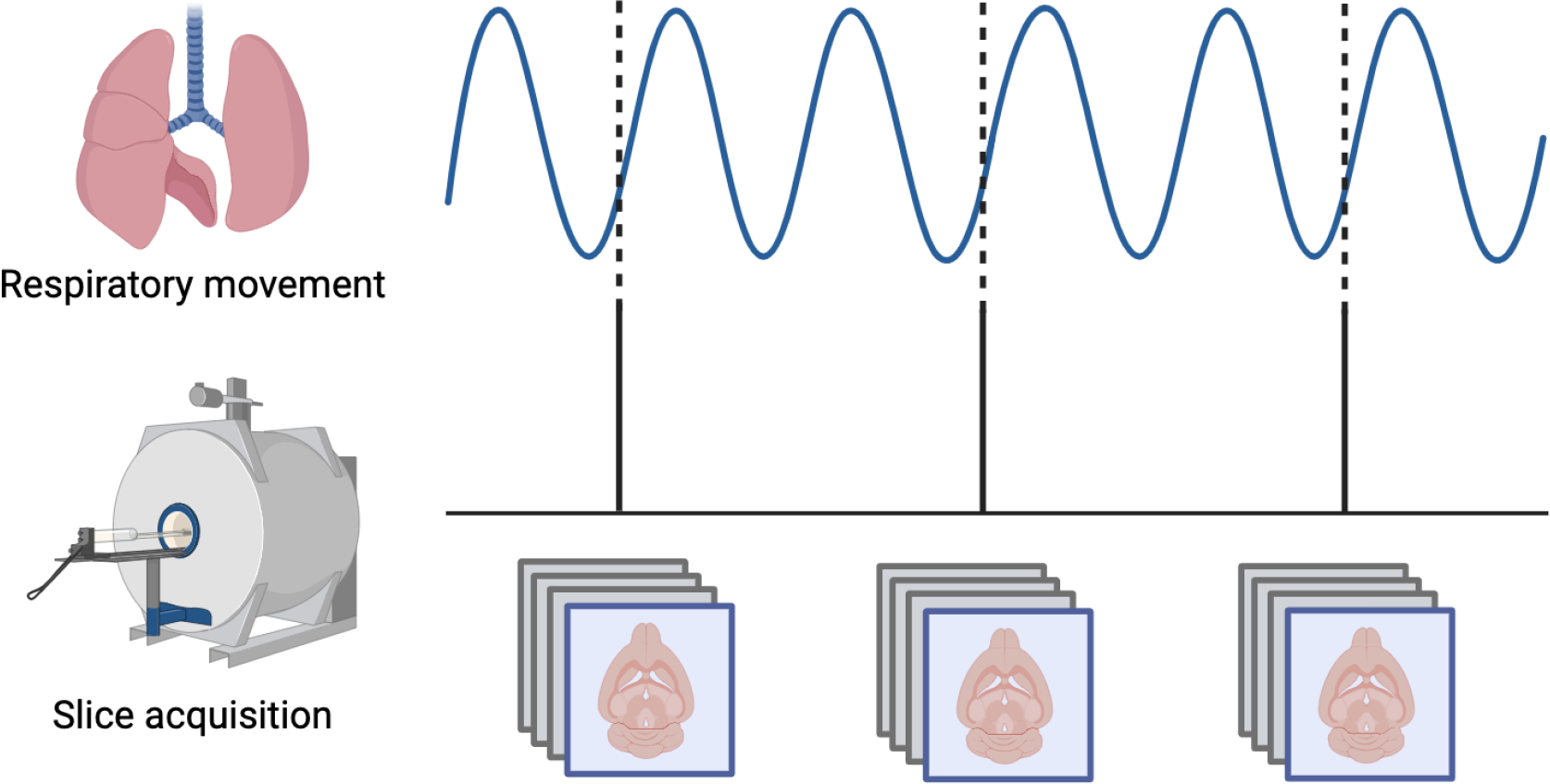
Mechanical ventilation in rats enables phase-locked acquisition of rs-fMRI data. The respiratory phase is continuously monitored and the interval between subsequent images is set so that each slice is always obtained at the same phase of the respiratory cycle, minimizing variability related to movement of the chest within the magnetic field.

### Preprocessing of rodent data

Preprocessing was performed using FSL ^35^, Analysis of Functional NeuroImages (AFNI) ^36^, ITK-SNAP ^37^, and ANTs ^38^. Distortion correction was applied to all scans using FSL Topup and registration to the 30^th^ volume of each scan was performed using AFNI 3dVolReg. Motion regression (6 parameters and up to 2 polynomials) and bandpass filtering (0.01 – 0.25 Hz) were performed in one step using AFNI 3dTProject. The bandpass range was chosen to accommodate all anesthetic conditions, as previous work has shown that BOLD is coherent with local field potentials for different frequencies under DMED (0.01 – 0.25 Hz) and ISO (0.01 – 0.1 Hz) ^39^. Spatial smoothing was not applied to this data. All scans were aligned to a single subject using direct, linear EPI to EPI registration via AFNI 3dAllineate and concatenated.

### Extraction and comparison of QPPs in rats

Data from all three anesthetic conditions were concatenated to enforce a common definition of QPPs. As in humans, QPPs were extracted using the complex PCA approach described in ^8^ with code available at (https://github.com/tsb46/complex_pca). Ten QPPs were obtained and varimax rotation was performed. The explained variance for each QPP and the magnitude and phase of each QPP at each time point were obtained. Based on the amount of variance explained, the top three QPPs were retained. The magnitude and phase of these QPPs were mapped, and movies that better demonstrate the evolution of activity over time were created. To investigate the occurrence of different components across scans and across rats, each time point was assigned a value (e.g., 1, 2, 3) based on the magnitude of the strongest QPP at that time point. Visual representations of the occurrence of states were created for all time points from each rat in each condition. The mean incidence of each state and the mean magnitude of each QPP for each condition were calculated. Repeated measures ANOVA was applied for each component to determine whether there was a significant effect of subject or condition.

## Results

### Distribution of QPPs in humans

The relative incidence of QPPs exhibited clear variability over time in HCP data. The number of subjects exhibiting QPP1 (i.e., global signal) at each time point increases at the expense of QPP2, while the incidence of QPP3 remains relatively stable (**Fig. 2**). When the QPPs are sorted for each subject rather than each time point, it is apparent that some subjects have relatively high ratios of QPP1 to QPP2, and others have much lower ratios. To further probe this finding, subjects were clustered into two groups with different patterns of QPP expression. For cluster 1, 41.8% of the time points were assigned to pattern 1, 22.7% to pattern 2, and 25.8% to pattern 3. For cluster 2, 23.9% of the time points were assigned to pattern 1, 21.6% to pattern 2, and 18.9% to pattern 3. When carpet plots are separated by cluster, it becomes clear that in cluster 1, QPP1 dominates and increases over the course of the scan at the expense of QPP2, while in cluster 2, time points are more evenly distributed across the three QPPs and maintain their relative representation over time (**Fig. 2**). QPP3 was more evenly distributed across clusters than QPPs 1 and 2. As a complementary analysis, the average magnitude of each QPP (rather than its relative incidence) across the first and last 200 time points of each scan was calculated across all subjects. The magnitude of each QPP was significantly greater (p<0.001) at the end of the scan than at the beginning of the scan, with QPP1 exhibiting the greatest change (**Fig. 2**). These results suggest that both the relative distribution of QPPs and their magnitudes provide information about changes in brain state that occur over the course of the scan.

**Figure 2.**
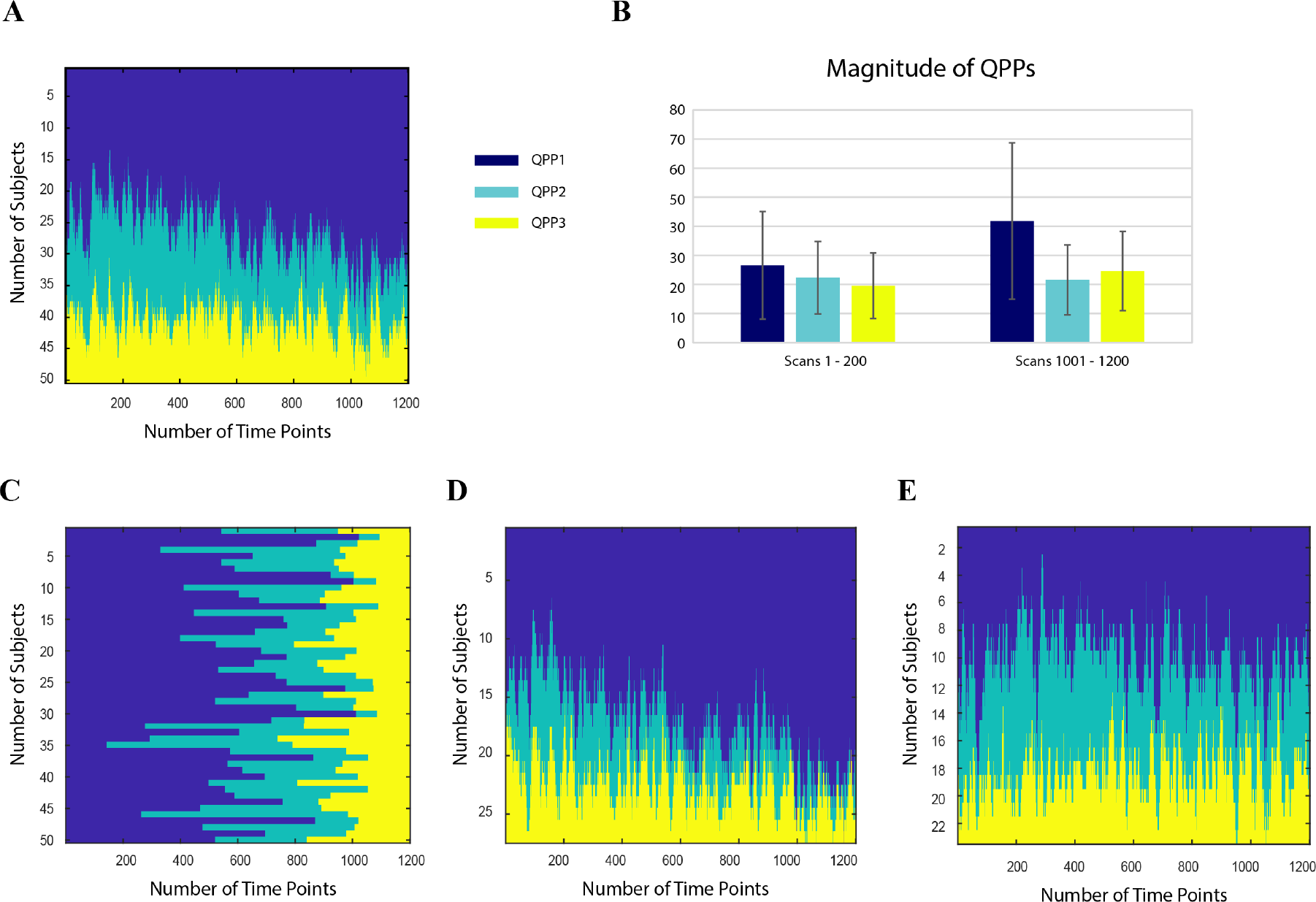
A) Distribution of QPPs 1 (dark blue), 2 (teal), and 3 (yellow) at each timepoint for all subjects. The incidence of state 1 increases over time and that of state 2 decreases. B) Average magnitude of each QPP in the beginning (first 200 time points) and end (last 200 time points) of the scan. The magnitude of each QPP was significantly greater (p<0.001) at the end of the scan than at the beginning of the scan. C) Distribution of QPPs for each subject over all time points. Differences in the relative incidence of different states can be observed. When subjects are clustered into two groups, one group (27 subjects) displays an even more prominent increase in QPP1 over time (D), while the distribution remains relatively constant in the second group (E; 23 subjects).

### Spatiotemporal patterns in ventilated rats with phase-locked acquisition

QPPs were detected across all anesthetic conditions. A plot of the variance explained by each component exhibited an ‘elbow’ around 3 QPPs (**Fig. 3**), similar to that observed in humans (Fig. 3 of ^8^). While the first component explains a larger percentage of the variation than the other two, the difference between the first component and the remaining components is less pronounced than in humans. Three view snapshots of the magnitude maps for each component are shown for the first 3 components in **Fig. 3** (movies available in the supplemental material).

**Figure 3.**
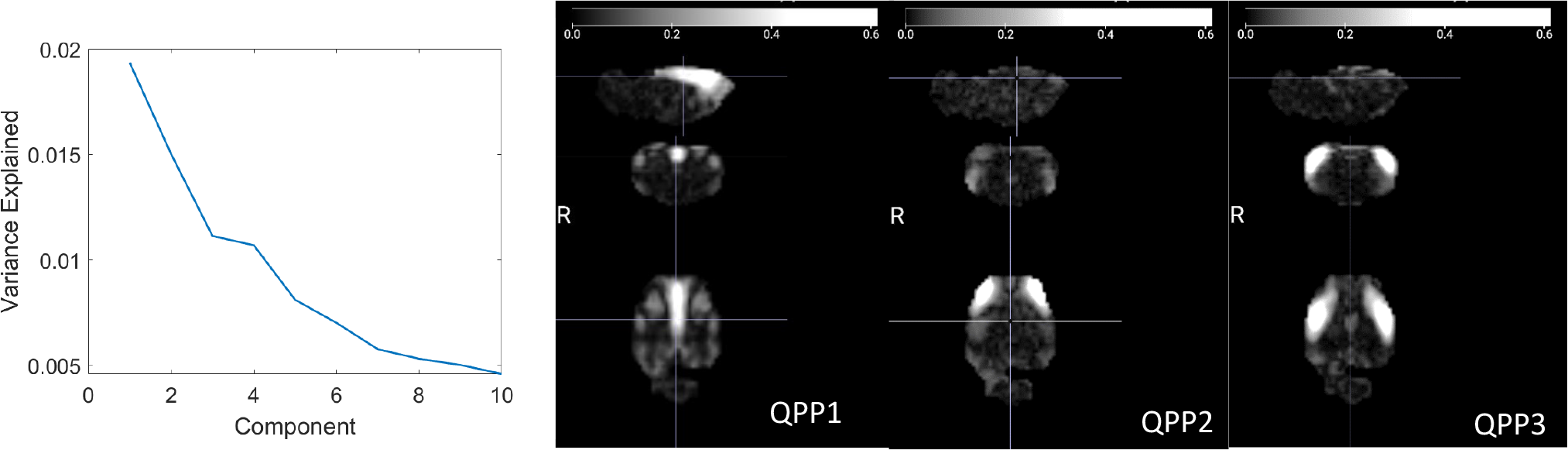
Left) Explained variance for each QPP. An elbow occurs around 3 QPPs, similar to prior work in humans. Right) Three directional snapshots of the whole-brain magnitude maps for the first 3 QPPs.

Prior work in rodents applied global signal regression and therefore only examined QPP2. As expected, QPP1 strongly resembles the pattern of global signal obtained from the same rats^14^, consistent with the observation that QPP1 captured the global signal in humans^8^. QPP2 and QPP3 both capture aspects of lateral-medial propagation from QPPs previously observed in rats after global signal regression^10,11^.

### Distribution of spatiotemporal patterns across animals and conditions

Animals exhibited variability in the expression of the spatiotemporal patterns over the course of the scan for each anesthetic condition (**Fig. 4**). No clear difference in the expression of patterns over the course of the scan was obtained, but unlike in humans where drowsiness is common as scans progress, the rats were expected to remain in a fairly stable state for each anesthetic condition. Carpet plots were not sorted across time or animal because the data acquisition periods varied substantially. The incidence of QPP1 was highest under ISO, while QPP2 was reduced in ISO compared to the other two conditions. To further explore the expression of different patterns under different anesthetic conditions, the average magnitude of each component was calculated. The repeated measures ANOVA found no significant effect of subject or anesthetic condition for the magnitude of any QPP, although magnitudes were lowest under ISO and highest under DMED. This suggests that the relative distribution rather than the magnitude of the various QPPs is most informative about brain state under different anesthetic conditions.

**Figure 4.**
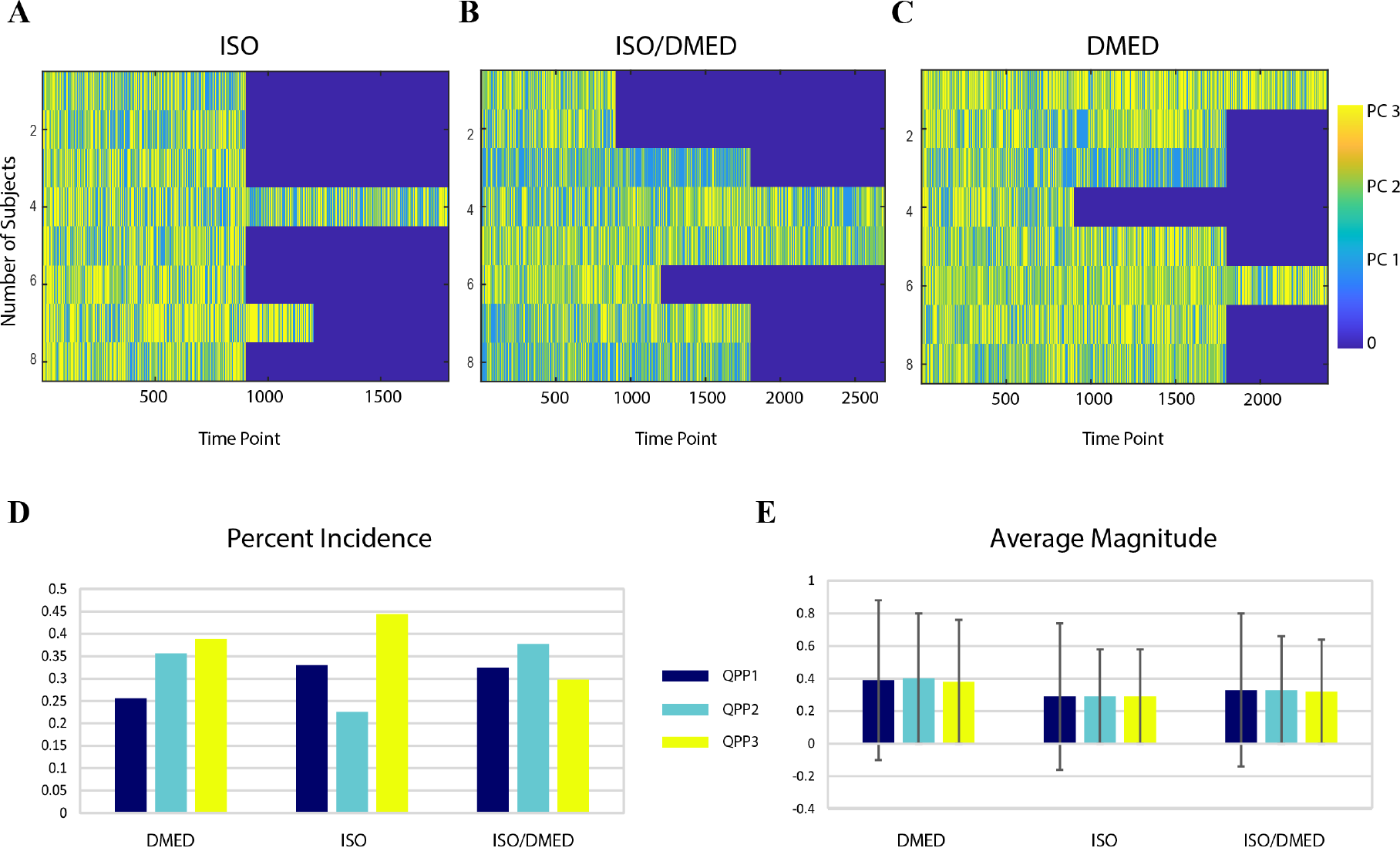
A) QPP state over time for each rat under different anesthetics (ISO, left; ISODMED, middle; DMED, right). Each row contains all time points from one rat. Substantial variation can be seen across animals and within a single animal over time. B) Overall percent incidence of each QPP for each of the three anesthetic conditions. Some QPPs occur more often in a single condition (e.g., 3 in ISO), or are reduced in particular anesthetic conditions (e.g., 1 for DMED). C) Average magnitude of each QPP in each condition. The magnitudes for ISO are lowest and for DMED are the highest.

## Discussion

Repeated whole-brain spatiotemporal patterns of intrinsic activity persist across a wide range of conditions and interact with transient tasks or stimuli. Based on the close links between global signal (QPP1) and arousal, and between QPP2, DMN/TPN anticorrelation, and attentional performance, we hypothesized that the relative distribution of the QPPs might reflect major changes in brain state. In humans, we observed an increase in QPP1 over the course of the scan at the expense of QPP2 that was most pronounced in a subgroup of subjects and which plausibly reflects a tendency toward increasing drowsiness over the course of the scan. In rodents, alterations in the distribution of the QPPs were observed based on the use of different anesthetics. Interestingly, in both humans and rodents, the state with presumably the lowest arousal level (i.e., the end rather than the beginning of the scan; ISO rather than DMED) has increased incidence of QPP1 relative to QPP2. This is consistent with the finding that the amplitude of the global signal increases with decreasing vigilance^18,27,28^, and that greater anticorrelation between the DMN and TPN (reflecting a stronger contribution from QPP2) is linked to better attentional performance^21,25^.

One of the key questions in the analysis of spatiotemporal patterns of activity is which features are most sensitive to changing conditions in the brain. The spatiotemporal patterns themselves are markedly similar across individuals and across scans^12^. We hypothesized that changes in brain state would be related to the relative distribution of the most prominent patterns, the magnitude of the patterns, or both. Our results suggest that both magnitude and incidence can be affected by changes in brain state. In humans, both the magnitude and incidence of QPP1 increase over the course of the scan, but in rats there is little differentiation in the magnitude of the QPPs across anesthetic conditions, and the relative incidence of the QPPs exhibits greater variability. Prior work has shown that the strength of the anticorrelation captured by QPP2 is weaker in subjects with ADHD compared to controls^30^. While a different algorithm was employed in that study, the results are consistent with a lower magnitude for QPP2 in ADHD. Together, these findings support the sensitivity of both magnitude and incidence of QPPs to changes in brain state.

### Source of QPPs

Despite the prominence of QPPs in rs-fMRI, little is known about the mechanisms that organize brain-wide activity into these few repeated spatiotemporal patterns. Multimodal studies have linked them to infraslow neural activity^40,41^, which is itself understudied, and QPPs exhibit reduced magnitude when slow waves are suppressed^42^. Structural connectivity undoubtedly plays a role in the organization of intrinsic brain activity, as shown by modeling studies that recreate much of the structure of functional connectivity by using a network representation of the brain’s structural connections along with a variety of neural mass models. In particular, a study in humans reproduces the division of the brain into two large, anticorrelated networks as observed in QPP2 in humans based on the structural connections and neural mass models^43^, although the patterns lack the complexity and propagation observed empirically. In rats, after surgical severance of the corpus callosum, QPPs continue to occur but their typical bilateral structure is disrupted^44^.

One potential source for brain-wide modulation is input from one or more deep brain neuromodulatory nuclei. For example, the basal forebrain the primary source of cholinergic projections to the cortex, and prior work has shown that unilateral stimulation of the NB alters GS (QPP1) ipsilaterally^45^. Other nuclei, such as the noradrenergic locus coeruleus or the serotonergic raphe nuclei, may also play a role. These systems are deeply interconnected and well-positioned to coordinate brain-wide patterns of activity. We have shown that QPPs distinguish between healthy mice and an Alzheimer’s model^46^; are altered in humans with ADHD compared to controls^30^; influence reaction time on a simple vigilance task^47^; are altered during task performance^13^; and interact with sensory stimuli^48^, all of which are consistent with a role for neuromodulatory input. Further experiments with chemogenetic or optogenetic manipulation of these nuclei^49^ may shed more light upon their role in the organization of QPPs.

### Global signal

The global signal is known to contain contributions from noise as well as widespread neural activity^50^. Temporal SNR can be difficult to interpret for BOLD fluctuations because multiple factors (some wanted, some unwanted) contribute to the amplitude of the fluctuations. In this case, we interpret the increased tSNR of the global signal as reflective of a greater contribution from neural activity relative to noise. Both the higher mean framewise displacement and the widespread increase in power in the global signal for the freely-breathing rats point to a higher noise level in the data.

In the mechanically-ventilated rats where global signal was reduced, the primary spatiotemporal pattern exhibited high amplitude along the midline, consistent with the regions most correlated to the global signal in ^14^. While this pattern explains the largest portion of the variance observed in the BOLD signal, the proportion is relatively less than in humans. We believe the primary explanation is the lack of smoothing in the rodent data compared to standard smoothing in the human data. Another possible explanation is that the signal in the ventilated rats includes fewer contributions from noise and from respiration in particular. This is evident from the lack of correlation between the global signal and nearby non-brain tissue described in ^14^. Our recent work in humans links the primary spatiotemporal pattern to changes in EEG power, pupil diameter, heart rate variability, respiratory volume, skin conductance, and peripheral vascular tone^51^, suggesting that the primary component in rats may also be linked to autonomic signaling. The animals are also anesthetized, which is likely to reduce the contribution of fluctuations in arousal.

### Relative occurrence across states

The three anesthetic states have all been used for rs-fMRI in rodents^7,14,52–54^, although most recently ISODMED has dominated the literature^52^. These anesthetics have a wide range of effects on neural activity (from enhancement of low frequency activity under DMED to cortex-wide spiking under deep ISO) and the vasculature (vasoconstriction under DMED, vasodilation under ISO). Interestingly, the relative incidence of QPPs 1 and 2 were reversed from ISO to DMED, possibly reflecting the relatively deep anesthesia obtained with ISO and the lighter sedation that occurs under DMED. Our results are consistent with our expectation that these different anesthetic regimens result in quite different brain states.

### Comparison to existing work in rodents

Prior studies in rodents had limited brain coverage and examined only a single QPP ^10,11,40,55,56^. In this study, we expand upon prior work to characterize multiple QPPs throughout the whole brain under three anesthetics expected to result in very different brain states. We show that mechanical ventilation phase-locked to image acquisition reduces apparent motion even in head-fixed, anesthetized rats where motion is minimal due to the reduced variability in disruption from the motion of the chest in the magnetic field. Under these ideal conditions, three spatiotemporal patterns of BOLD fluctuations explain much of the variance in the signal, as in humans, across anesthetic conditions. QPPs have been observed in species from mice to humans^9–13,13,46,57^, which is not surprising given that the functional networks obtained based on the patterns of intrinsic activity are also present across species^4^. Thus, the clear differences in the magnitude and incidence of the patterns that demonstrate their sensitivity to induced changes in whole brain activity suggest that the relative expression of different patterns may provide information about the changes in brain state related to nonlocalized processes such as autonomic processing, arousal, emotion, and more across species.

### Limitations

In the data from humans, no simultaneous measure of arousal (e.g., EEG) was available. Nevertheless, it has been shown that drowsiness tends to increase over the course of the scan^58^, and our results are consistent with these findings. The assignment of each time point to a single dominant QPP (1, 2, or 3) provides a simplistic view of the current brain state. While well-suited to the current investigation of changes in brain state, a more nuanced definition based on the relative magnitudes of all QPPs might provide further insight.

In the rodents, while phase-locked acquisition reduces noises, it has drawbacks in that it requires more invasive preparation (intubation) and reduces flexibility in the choice of TR, which must be a multiple of the respiratory cycle. It may also have more subtle drawbacks related to alterations arising from the lack of top-down control, although bottom-up feedback is maintained ^59^.

For the anesthesia study in rats, data from all conditions was concatenated. A preliminary analysis run on separate data for each condition suggested that the spatiotemporal patterns extracted were similar, supporting concatenation. The observation that all three patterns are observed in all three states also suggests that concatenation is appropriate. Further work should investigate more subtle differences in the spatiotemporal patterns and timing across groups.

## Acknowledgements and Funding

Emory’s Center for Systems Imaging Core, 1R01 MH111416, 1R01NS078095, 1R01AG062581.

