## Supplemental Methods for "Variation in the Distribution of Large-scale Spatiotemporal Patterns of Activity Across Brain States"

*Free breathing vs. ventilation with phase-locked image acquisition in rats*. For each rat, anesthesia was induced using 5% isoflurane (ISO), and was maintained at 2% ISO during setup. Animals were placed in a homemade custom MRI cradle and positioned with their teeth secured in a bite bar and their heads fixed using ear bars. Physiological parameter (oxygen saturation, temperature, heart rate, and CO2 (for intubated scans)) were monitored and recorded throughout the experiment.

To compare free breathing and mechanical ventilation, four Sprague-Dawley rats (2 male and 2 female rats, 230- 400g) were imaged under both conditions. Free breathing scans were collected under ~0.5% ISO and dexmedetomidine (DMED), a combination widely used for functional studies^30^. The isoflurane was delivered in a mixture of 30% O2 and 70% air continuously through the nosecone. DMED (0.5 mL/kg) was injected subcutaneously prior to the scan and a subcutaneous infusion line of DMED was set up for the duration of the scan (0.017 mL/kg/hr). Each animal was given a week to recover prior to being intubated. Intubated scans used the same anesthetic doses as the free breathing scan with the addition of a second infusion line for the paralytic pancuronium (1.5mg/kg/hr). Mechanical ventilation was performed at approximately 1 Hz. For the comparison between freely-breathing and mechanically-ventilated animals, each rs-fMRI scan was 30 minutes long.

All rodent imaging was performed on a 9.4T/20 cm horizontal bore small animal MRI system with a homemade transmit/receive surface coil ~2 cm in diameter. Anatomical scans were obtained using a T2-weighted RARE sequence (TR = 3500 ms, TE = 11 ms, 24 axial slices, 0.5 mm^3^ isotropic voxels). All rs-fMRI scans were acquired using a gradient-echo echo-planar imaging (EPI) sequence with field of view (FOV) 35mm x 35 mm, matrix size 70x70, and 24 axial slices for whole-brain coverage, resulting in isotropic voxels of 500 microns. Acquisition parameters include flip angle = 68.4°, bandwidth = 216.45 kHz, TE = 15 ms, TR = 1250 ms, with partial Fourier encoding (encoding factor 1.4) to reduce the length of the echo train. A 3-volume reversed blip EPI image with the same parameters was acquired before each longer functional scan for TOPUP correction^30,31^. All functional EPI scans included saturation bands to minimize signal from frontal and ventral regions outside the brain and were preceded by 10 dummy scans to ensure the signal reached steady state.

All rs-fMRI scans for mechanically-ventilated rats were phase-locked, meaning that the time between subsequent images was set to a multiple of the animal’s respiratory rate so that each image was acquired during the same phase of the respiratory cycle^14^ (**Fig. 1**). This practice was adopted here to limit any effects of motion that could arise due to movement of the chest cavity and volumes being imaged at different points in the respiratory cycle.

Preprocessing was performed using inhouse MATLAB scipts that work with SPM12 (https://www.fil.ion.ucl.ac.uk/spm/software/spm12/). Preprocessing steps include slice time correction, motion correction and smoothing using spm_slice_timing, spm_realign, spm_reslice and spm_smooth. Brain masking was done manually in FSL and final data was bandpass filtered between 0.01-0.25Hz.

To assess the minimization of physiological noise in the intubated condition the frame wise displacement, temporal signal to noise ratio (tSNR) in the global signal and the power spectrum of the global signal were analyzed. Analysis for all 4 rats was done individually. All 6 motion parameters for each scan were used to calculate the framewise displacement (FD) which is the sum of the absolute temporal derivatives of the 3 translation and 3 rotational parameters. The FD was calculated for each 30-minute scan and then averaged across the two conditions. The global signal (GS) was calculated by taking the mean of the voxel time series within the brain, the first and last 2 minutes of the scan were disregarded. Removal of the first couple TRs is commonly done to eliminate magnetization artifact. The tSNR was calculated by dividing the mean by the standard deviation for each time course. The power spectrum for each GS time series was then obtained using the MATLAB pspectrum function. Here as well the analysis was performed on a per scan basis and then averaged to obtain results for the two conditions. The light-colored lines on the power spectrum plot represent the standard error for each condition.

*Comparison of freely-breathing to intubated animals*. Mean framewise displacement was lower in intubated rats compared to freely breathing rats (Fig. S1B; 0.0388 ± 0.0075 free breathing, 0.0304 ± 0.0041 intubated). Because both groups of animals were anesthetized and head-fixed, the reduction in motion is most likely attributable to the reduction of variation in chest position during image acquisition rather than any overt motion of the animal. Note that the use of a surface coil for transmission and reception of the RF signal already limits the contribution from areas far from the brain, and the noise from respiration in freely-breathing rats is likely to be even more pronounced when volume coils are used.

The global signal has been shown to contain both contributions from noise and from widespread neural activity. In intubated rats, the tSNR of the global signal (Fig. S1C; 0.01 ± 0.0077) is higher than in freely-breathing rats (0.0017 ± 0.0013). Together with the reduction in motion, this suggests that the relative contribution of noise vs. neural activity is reduced in the phase-locked acquisition. Examination of the power spectra for each condition indicates that power in the low frequencies is higher in freely-breathing animals, particularly below 0.05 Hz but with clear differences up to ~0.1 Hz (Fig. S1D).

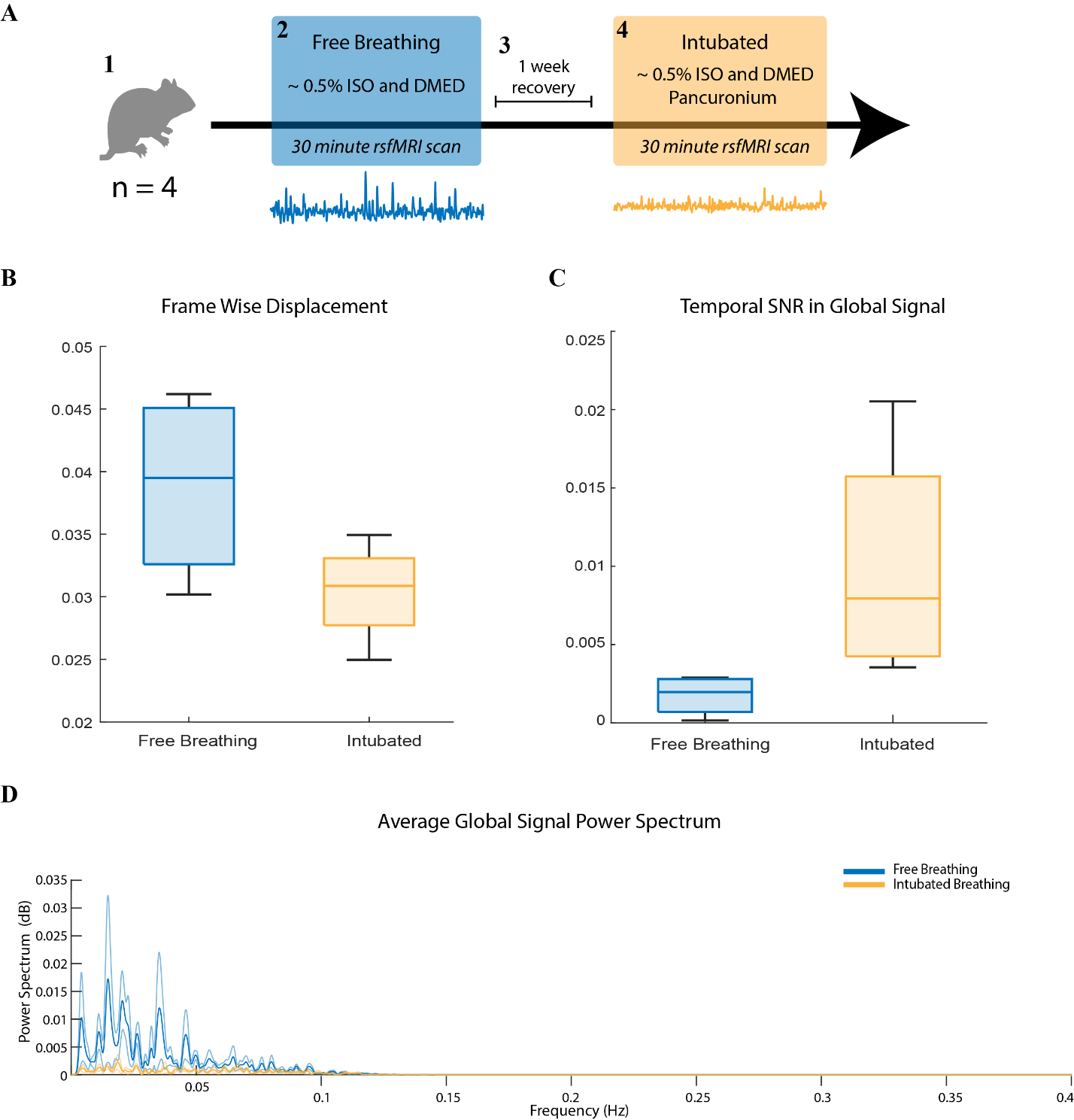

*Figure S1. A) Experimental design for comparison between freely-breathing and phase-locked ventilation conditions. B) Mean framewise displacement in free breathing (mean = 0.0388, std = 0.0075) vs. intubated rats (mean = 0.0304, std = 0.0041). C) Temporal SNR of the global signal is higher in intubated rats (mean = 0.01, std = 0.0077) than in freely-breathing animals (mean = 0.0017, std = 0.0013). D) Global signal power spectra show higher fluctuations in the low frequencies in freely-breathing animals, particularly below 0.05 Hz.*

**Movies S2-S4**. Videos of QPPs 1-3.
